# iGIST - a kinetic bioassay for pertussis toxin based on its effect on inhibitory GPCR signaling

**DOI:** 10.1101/2020.09.29.318451

**Authors:** Valeriy M. Paramonov, Cecilia Sahlgren, Adolfo Rivero-Müller, Arto T. Pulliainen

## Abstract

Detection of pertussis toxin (PTX) activity is instrumental for the development and manufacturing of pertussis vaccines. These quality and safety measures require annually thousands of mice. Here, we describe iGIST (<u>I</u>nterference in <u>G</u>αi-mediated <u>S</u>ignal <u>T</u>ransduction) - an animal-free kinetic bioassay for detection of PTX by measuring its effect on inhibitory G protein-coupled receptor (GPCR) signaling. PTX ADP-ribosylates inhibitory α-subunits of the heterotrimeric G proteins, thereby perturbing the inhibitory GPCR signaling. iGIST is based on HEK293 cells co-expressing a somatostatin receptor 2 (SSTR2), which is an inhibitory GPCR controllable by a high affinity agonist octreotide, and a luminescent 3’5’-cyclic adenosine monophosphate (cAMP) probe. iGIST has a low sensitivity threshold in picogram/ml range of PTX, surpassing by 100-fold in a parallel analysis the currently used *in vitro* end-point technique to detect PTX, the cluster formation assay (CFA) in Chinese hamster ovary cells. iGIST also detects PTX in complex samples, i.e. a commercial PTX- toxoid containing pertussis vaccine that was spiked with an active PTX. iGIST has an objective digital readout and is observer-independent, offering prospects for automation. iGIST emerges as a promising animal-free alternative to detect PTX activity in the development and manufacturing of pertussis vaccines. iGIST is also expected to facilitate basic PTX research, including identification and characterization of novel compounds interfering with PTX.

---

The Gram-negative bacterium *Bordetella pertussis* is the etiological agent of the whooping cough, i.e. pertussis. Whooping cough is a globally distributed acute respiratory disease, affecting all age groups ^1^. However, infants and young children comprise the highest risk cohort, where the disease may lead to death despite hospital intensive care and use of antibiotics ^1^. Despite the global vaccine campaign pertussis remains endemic, causing outbreaks in many regions of the world, and the disease incidence is increasing ^2^. Moreover, macrolide resistant *B. pertussis* strains have been reported ^3, 4^. The data highlights the need to improve the current vaccine formulations and vaccination campaigns.

Pertussis toxin (PTX) is the major virulence factor of *B. pertussis* ^5^, a protein complex secreted from the bacteria via the Sec-pathway and the Ptl type IV secretion system ^6^. PTX is composed of five non-covalently bound subunits (PtxS1-S5), which are arranged in an AB_5_-topology ^7, 8^. The B_5_ oligomer is formed by the PtxS2-S5 (PtxS2, PtxS3, PtxS5 and 2 copies of PtxS4) ^7, 8^ and mediates binding of the secreted AB_5_ holotoxin on the host cell surface in a carbohydrate-dependent manner ^8^. Subsequent cell entry is followed by dissociation of the B_5_ oligomer and the PtxS1 ^9^. The liberated PtxS1, which belongs to the family of ADP-ribosyltransferases ^10^, ADP-ribosylates a single C-terminal cysteine residue in inhibitory α-subunits of most heterotrimeric (αβγ) G protein superfamily members, such as Gαi, Gαo, and Gαt ^11-13^. The resulting bulky ADP-ribose modification disrupts inhibitory α-subunit interaction with G protein-coupled receptors (GPCRs), preventing formation of the Gαβγ-GPCR complex and thereby perturbing GPCR agonist-induced signaling ^14, 15^. Although the pathogenic manifestations are still a matter of debate ^5^, one well-recognized molecular downstream effect is an altered 3’5’-cyclic adenosine monophosphate (cAMP) signaling ^16^. This is attributed to the diminished inhibitory control of PTX-modified Gαi on the cAMP-producing adenylyl cyclases (ACs) (**Table of Contents Graphic**).

A detoxified form of PTX (PTX-toxoid) is a core component of the pertussis acellular vaccines (ACVs), where it is typically included at µg/ml levels. Thus, tests for residual PTX activity are instrumental for ACV development and manufacturing. Mouse histamine sensitization test (HIST) is a benchmark PTX assay in the vaccine industry capable of detecting PTX at ng/ml levels ^17^. HIST is based on the early observation of PTX-treated mice becoming sensitive to histamine ^18^. Mice are exposed to PTX-containing preparations, challenged with histamine and monitored for death ^17^. Though it has a long record in the industry, HIST is a terminal assay causing profound stress for the animals. Besides, HIST requires large amounts of animals, with a recent global annual estimate of 65.000 mice ^17^.

The most widely debated animal-free alternative to HIST builds on the early findings by Hewlett et al., who observed phenotypic alterations, described as cell rounding and cell cluster formation, in Chinese Hamster Ovary (CHO) cells exposed to PTX ^19^ (**Supplementary Videos 1-2**). The resulting test, designated as a cluster formation assay (CFA), is based on visual grading of the cell clustering in CHO cell monolayers upon PTX treatment and can detect ng/ml levels of PTX ^17^. However, the CFA is an observer-dependent end-point test, suffering from subjectivity bias and considerable inter-assay variability ^17^. Also, the molecular basis of the PTX-evoked clustering in CHO cells, similar to the mechanism of histamine hypersensitivity in HIST, remain poorly understood. Despite these limitations, the European Pharmacopoeia Commission has decided that CFA can be used instead of HIST for safety assurance of the currently marketed pertussis ACVs ^17^, based largely on work of Isbrucker et al. ^20^, effective as of January 2020. Recently, Biological Reference Preparation batch 1 (BRP1) of PTX was introduced in order to control the inter-assay variability of CFA ^21^.

Improved alternatives to CFA, based on the mechanistic understanding of PTX cellular effects, have been actively sought for ^17^. Available biochemical assays for PTX measure either the PtxS1-catalyzed ADP-ribosylation of a C-terminal peptide of Gαi with HPLC ^22^ or binding of the pentameric PtxS2-S5 oligomer to carbohydrate structures with ELISA ^23^. Both the assays have objective readouts, but capture only distinct PTX activities under artificial cell-free *in vitro* conditions. DNA microarrays have been utilized to identify PTX-induced gene expression signatures either in rat tissues ^24, 25^ or in *in vitro* cultured human cells ^26^. Practical applications have not yet emerged from these studies. Hoonakker et al. exposed rat vascular smooth muscle cells (A10 cells) to PTX and determined the amount of cAMP in cell lysates with an end point ELISA ^27^. PTX did not increase the amount of cAMP when incubated alone with the cells, but it potentiated isoproterenol-induced elevation of cAMP ^27^. Isoproterenol binds to β-adrenergic receptors ^28^, which leads to activation of Gαs and thereby to subsequent stimulation of the cAMP-producing ACs. In an extension of their work, Hoonakker et al. detected PTX effects in A10 and CHO cells with a cAMP response element (CRE)-driven luciferase reporter ^29^. In agreement with their earlier cAMP ELISA study ^27^, PTX did not increase the CRE-reporter activity by itself, but it did enhance cAMP responses to isoproterenol or forskolin (FSK) ^29^. FSK activates ACs by intercalating the C1 and C2 subunits of ACs into the catalytically active cAMP-producing form ^30^. Although the detailed molecular basis of the CRE-reporter assay was not reported, the PTX-mediated blockage of basally active Gαi signaling was probably sufficient to allow enhanced cAMP accumulation upon pharmacological AC stimulation. The CRE-reporter assay has a low ng/ml-range sensitivity for PTX ^29^, comparable to CFA ^19^, yet the question of its practical use in vaccine industry awaits further studies.

In this work, we set out to establish a sensitive microtiter plate format bioassay for PTX, based on kinetic measurement of intracellular cAMP levels in living cells in combination with a defined and tightly controllable inhibitory GPCR pathway.

## EXPERIMENTAL SECTION

### Compounds and reagents

All the reagents were dissolved in ultrapure water (Milli-Q; resistivity >18mΩ*cm), if not specified otherwise. PTX preps were obtained from List Biological Labs (#179A, Lot#179216A2A, aka PTX_#1_, stock of 200 μg/ml in 50 mM Tris, 10 mM glycine, 0.5 M NaCl, 50% (v/v) glycerol in H_2_O, pH 7.5; kept aliquoted at -20 °C) and Invitrogen (#PHZ1174, Lot#75356597A, aka PTX_#2,_ stock of 100 μg/ml in 10 mM Na_2_HPO_4_ and 50 mM NaCl in H_2_O; kept aliquoted at +4 °C). Control solvents for both the PTX preps (SolC_#1_ and SolC_#2_), of chemical composition identical to the specified above, were prepared in-house, filter-sterilized and kept at +4 °C. Octreotide acetate was obtained from Bachem (#H-5972) and kept at -80°C as single-use 100 μM aliquots. FSK was from LC laboratories (#F-9929) and kept aliquoted (10 mM) in dimethyl sulfoxide (DMSO) at -20°C. Boostrix vaccine was from GlaxoSmithKline (tetanus toxoid, reduced diphtheria toxoid and acellular pertussis vaccine, adsorbed; lot# AC37B272AK; 16 μg/ml of formaldehyde and glutaraldehyde-inactivated PTX in 9 mg/ml NaCl with ≤ 0.78 mg/ml of Al as aluminum hydroxide and ≤ 200 μg/ml Tween80; full composition – as described by the manufacturer).

### Cell lines

Human embryonic kidney cell line (HEK293) was obtained from American Type Culture Collection (ATCC, #CRL-1573). HEK293 with stable overexpression of Gs22/cAMP probe, as well as the derived sensor cells with stable overexpression of SSTR2 (aka HEK-Gs/SSTR2_HA), were developed and characterized by us earlier ^31-33^. Chinese hamster ovary cells (CHO) were either recovered from the local cell line repository of the Institute of Biomedicine, University of Turku, Finland (liquid N_2_ storage – a cryovial of the stock culture of 1998; aka CHO_#1_) or provided as a kind gift by Dr Aylin C. Hanyaloglu (Institute of Reproductive and Developmental Biology, Imperial College London, UK; aka CHO_#2_). HEK293 and CHO cells were cultured in Dulbecco’s Modified Eagle Medium/Nutrient Mixture F-12 (DMEM/F-12; Gibco, #11320033), supplemented with 10% (w/v) of heat-inactivated fetal bovine serum (iFBS; Biowest, #S1810), in the incubator conditions (+37°C in humidified atmosphere with 5% CO2). Only verified mycoplasma-negative cells were used for the experiments. Cell counts were performed with TC20 automated cell counter (Bio-Rad Labs).

### iGIST bioassay for PTX activity

The sensor cells were seeded on the day of the experiment into tissue culture-treated polystyrene 96-well plates with light-tight walls and translucent bottom (ViewPlate-96, PerkinElmer, Cat#6005181) as 60,000 cells per well in 180 μl of complete medium, and incubated for 4-6 h (+37°C in humidified atmosphere with 5% CO_2_) to allow for attachment. Next, the freshly prepared PTX dilutions or matched SolC dilutions (both in 25 mM HEPES, pH 7.4) were added to the wells as 20 μl of 10x solutions to yield the desired 1x working concentration. Non-treated controls received 20 μl of the specified HEPES (25 mM, pH 7.4) buffer per well. Further, the plates were placed back in the incubator and kept at the above-specified conditions for the scheduled time to allow for PTX to act. Once the desired exposure time had elapsed, the plates were retrieved from the incubator, the medium was removed and the wells were refilled with 45 μl of the freshly prepared inducing medium (IndMed), comprised of 2% (v/v) of GloSensor reagent (Promega, #E1290; corresponds to the final working concentration of 0.612 mg/ml, with the original stock of 30,6 mg/ml in 10 mM HEPES, pH 7.5) and 200 μM of a non-selective family-wide phosphodiesterase inhibitor 3-Isobutyl-1-methylxanthine (IBMX; Sigma, #I5879) in a mix of DMEM/F-12 medium (50/50, v/v) and CO_2_-independent medium (Gibco, #18045-054; 4v of DMEM/F12 per 5v of CO_2_-independent medium), supplemented with 0.1% (w/v) of bovine serum albumin (BSA). After equilibration for 45 min at RT in the dark, the plate was inserted into a microtiter plate reader (EnSight, PerkinElmer) and the light output, denoted as a baseline signal, was captured for 15-20 min at RT. Next, the plate was removed from the reader and the wells were spiked with either 5 μl of freshly prepared solutions, having all the desired components at 10x of the final concentration in 25 mM HEPES, pH 7.4, or 5 μl of respective controls [i.e., HEPES solutions of either 100 nM Oct or 1% (v/v) DMSO, or just HEPES buffer]. Final concentrations of FSK and Oct in the assay equaled 10 μM and 10 nM, if not specified otherwise. As the 10 mM FSK stock was in DMSO, final DMSO concentration in all FSK-spiked samples and DMSO controls equaled 0.1% (v/v). After spiking, the plate was immediately re-inserted into the reader and the luminescence, now denoted as induced signal, was further recorded for the time required (typically, for 45-60 min). The described assay conditions [i.e., at RT, IndMed with 2% (v/v) of GloSensor reagent and 200 μM of IBMX, stimulation with 10 μM of FSK] are referred to as standard throughout the text. The assay with Boostrix vaccine followed the same design. First, 20 µl of 10x of vaccine with or without external PTX_#1_ at a fixed concentration of 1000 ng/ml in Milli-Q H_2_O was added to the sensor cells in 180 µl/well of the complete medium, yielding the final desired 1x of vaccine dilution (dilution range 1:10 - 1:10^−6^) +/- 100 ng/ml PTX_#1_. Next, the sensor cells were incubated for 24 h (+37°C in humidified atmosphere with 5% CO_2_) before exposure to 100 nM of Oct and 10 µM of FSK.

For initial inspection and qualitative analyses of iGIST data, the captured luminescent reads were plotted as intracellular cAMP kinetic curves (luminescence *vs* time) and subjected to visual assessment. Subsequent quantitative analyses involved several steps of data transformation and were carried out as follows. Firstly, cAMP kinetic curves were processed to obtain baseline signal-subtracted Area Under the Curve (AUC)-values by subtracting the average baseline signal from the AUC-value for the period of induced signal. This was done either with the corresponding operator of GraphPad Prism software or via a custom-written script, both employing the trapezoidal rule ^34^ and producing similar results. The obtained AUC-values were further divided by the average AUC-value of FSK response in control cells (control-AUC_FSK_), i.e. sensor cells spiked only with FSK after the baseline signal capture. This yielded FSK-normalized AUC%-values (PTX-AUC% or SolC-AUC%). Finally, the PTX-AUC%- and SolC-AUC%-values for FSK *vs* FSK + Oct 10 nM responses were combined to obtain the following two ratiometric values: i) Gαi signal relay index [Gαi-SRI; separately calculated for PTX and SolC as AUC% _FSK_ / AUC% _FSK + Oct 10 nM_], and ii) comparative Gαi signal relay index [comparative Gαi-SRI; calculated as a ratio of AUC% _FSK_ / AUC% _FSK + Oct 10nM_ for PTX to AUC% _FSK_ / AUC% _FSK + Oct 10 nM_ for SolC]. At full abrogation of Gαi signaling by PTX, the sensor cells are expected to completely lose responsiveness to Oct, with Gαi-SRI approaching 1.0. Gαi-SRI allows to separately estimate dose effects of PTX and SolC on Gαi signaling, whilst the derived comparative Gαi-SRI integrates the effects of matched PTX and SolC doses into a single numerical value. Comparative Gαi-SRI thus accounts for any solvent effects and reveals the genuine solvent-corrected effect of PTX. Further details on luminescence data processing are covered in our earlier work ^32^. Schematics of iGIST output values and of their calculations is shown in **Figure S1**.

### CHO cluster formation assay and confluence analysis

Cluster formation assay (CFA) was carried out based on the original descriptions by Hewlett et al. ^19^, with the following modifications. CHO cells were seeded into flat-bottom 96-well plates (#655180; Greiner) as 10,000 cells/well in 180 μl of the complete medium. The plates were then placed in the incubator for 4-6 hours to allow for cell attachment. Further, the cells were treated with 20 μl/well of PTX or matched SolC, as specified in the previous section. Next, the plates were inserted into IncuCyte HD live cell imager (Essen BioScience), integrated with the cell culture incubator (+37°C, humidified atmosphere with 5% CO_2_), and immediately subjected to continuous phase-contrast imaging (1 snapshot every 30-60 min, up to 72h from the moment of treatment initiation). As the culture plates were placed into the imager within 5 min of treatment, the first imaging time point was also taken for the time point 0 in terms of the subsequent image analysis. No medium exchange or other perturbations were performed during the imaging.

Visual grading of morphological changes in CHO cells, exposed to different doses of PTX or matched SolC, were performed by six independent observers (2 males, 4 females; all adults), who had never dealt with such type of analysis earlier. After a short introductory tutorial (a single parallel session with all the observers) on how the grading is expected to be implemented, including review of selected examples of morphological changes in CHO cells in response to varying doses of PTX/SolC, the observers received the identical set of phase-contrast images of CHO cells, assembled as slides of 3x images each [two different filed of views (FoVs) of CHO cells in different wells of the same 96-well plate, exposed to the same dose of PTX (i.e. two technical replicates) *vs* one FoV of the cells that received the matched level of SolC in the same experiment]. The observers remained blinded to the PTX dose, exposure time and CHO strain information (i.e., CHO_#1_ *vs* CHO_#2_), but were aware of the nature of treatments on every slide (i.e., FoVs for PTX and SolC were explicitly labeled). The grading followed a simple 3-tier scale (0 - no effect; 1 - equivocal response; 2 - clear response) and relied on visual comparison of FoV for PTX samples with the FoV of matched SolC by every observer. The resulting grades were entered into spreadsheets, available from the authors upon request, and processed to yield the average grades (out of 6 observers; +/-SD) for every PTX dose/exposure in a given CHO strain in a given experiment. The resulting averages were eventually used to compute the respective final mean grades (with SEM and 90% CI) across several independent experiments. The final grade of 1.5 was selected for an arbitrary cut-off of a clear response. Confluence analyses for CHO cells, reflective of the surface occupancy by cells in a given FoV (with 100% corresponding to the full confluence, i.e. when a FoV is fully covered with cells), were performed with IncuCyte software (build 2010A Rev3; Confluence v.1.5 operator) on the same phase-contrast image sets that were utilized for visual grading.

### Data transformation, curve fitting and statistics

Data transformations, CI calculation and inferential statistics were carried out with GraphPad Prism v8.4.1 package (GraphPad Software). Dose-response curve fitting [log (inhibitor) versus response - variable slope (Y=Bottom + (Top-Bottom)/(1+10^((LogIC_50_-X)*Hill Slope)) for non-linear regression and «Y=B0 + B1*X + B2*X^2 + B3*X^3 + B4*X^4 + B5*X^5» for 5^th^ order polynomial regression] was performed using the respective operators of GraphPad Prism software. Comparisons of PTX *vs* SolC dose effects were performed with either a paired ratio two-tailed t test or a simple two-tailed t test (for the effects, expressed as AUC%-values or through a Gαi-SRI, respectively). Level of significance was set to <0.1 for all of the tests (on the figures, one (*), two (**), three (***) and four (****) asterisks indicate p values in the following ranges - [0.05;0.1), [0.01;0.05), [0.001;0.01) and <0.001, respectively).

## RESULTS AND DISCUSSION

### iGIST bioassay robustly detects PTX-induced abrogation of Gαi signaling

iGIST is based on stably-transfected HEK293 sensor cells (HEK-Gs/SSTR2_HA), co-expressing somatostatin receptor 2 (SSTR2), which is a well-characterized GPCR negatively regulating the cAMP-producing ACs through PTX-targeted Gαi, and a luminescent cAMP probe GloSensor-22F ^35, 36^. GloSensor-22F, originally introduced by Wood et al. ^35, 36^, represents a cAMP-binding domain of protein kinase A fused to a circularly permuted *Photinus pyralis* luciferase, jointly functioning as a sensitive and reversible cAMP probe in living cells. The sensor cells were earlier established in-house in HEK293 background, which has low endogenous expression of SSTR2 ^32, 33^, and used to measure SSTR2-mediated signaling upon exposure to various ligands. In iGIST, activities of SSRT2 and ACs are controlled with a high-affinity synthetic peptide agonist octreotide (Oct) ^37^ and forskolin (FSK), respectively. Oct induces potent and dose-dependent activation of SSTR2 with an IC_50_ of 0.3 nM (in iGIST typically used at 10 nM) ^32^. FSK activates ACs by intercalating the C1 and C2 subunits into the catalytically active form ^30^, which readily boosts intracellular cAMP levels and facilitates registration of counter-acting stimuli, i.e. inhibition of ACs via the Oct/SSTR2-induced Gαi signaling. The PTX-catalyzed ADP-ribosylation of Gαi prevents Gαi-GPCR coupling with ensuing loss of Gαi-mediated inhibitory control on ACs ^11-13^. Schematics of the molecular basis of iGIST bioassay is shown in **Figure 1**. To the best of our knowledge, no literature exists on the physiological role of SSTR2 in whooping cough. In principle, iGIST could be based on alternative iGPCRs, as long as they can be efficiently expressed and pharmacologically stimulated in sensor cells.

**Figure 1.**
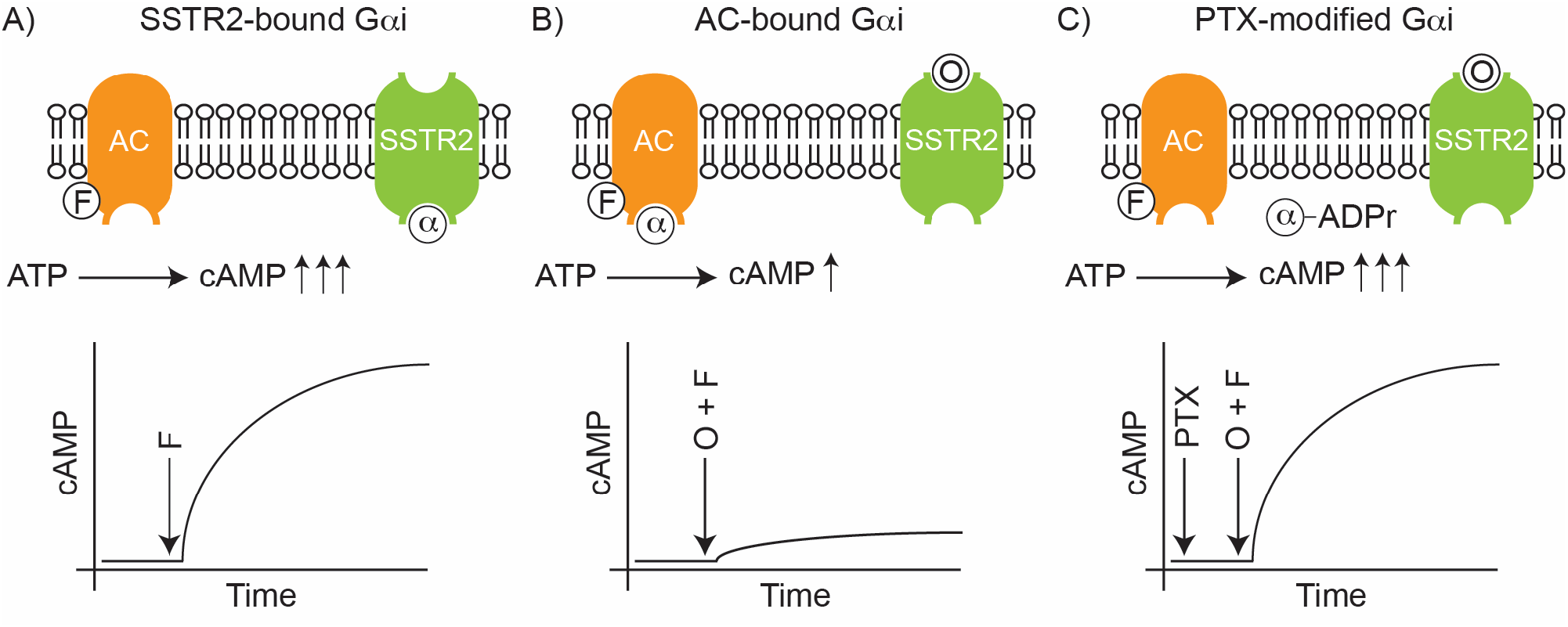
Schematics of the molecular basis of iGIST bioassay to detect PTX. iGIST is a living cell-based bioassay for PTX, measuring the PTX-induced alterations in Gαi signaling in HEK293 cells, stably transfected with Gαi-coupled SSTR2 GPCR and luminescent cAMP probe GloSensor-22F. **A)** Forskolin (F) binds to and activates cAMP-producing adenylyl cyclase (AC), leading to increased generation of intracellular cAMP. **B)** Octreotide (O) is a high affinity agonist of SSTR2. Binding of octreotide to SSTR2 activates AC-inhibitory Gαi protein (α), which binds to ACs and counteracts forskolin-induced generation of cAMP. **C)** PTX ADP-ribosylates the AC-inhibitory Gαi protein (α), thereby preventing Gαi-SSTR2 interaction. Thus, SSTR2 cannot inhibit ACs through Gαi any longer, and the forskolin-induced cAMP generation rate is restored.

We incubated the sensor cells with a commercial PTX_#1_ preparation and subsequently challenged them with FSK or FSK + Oct 10 nM. Importantly, in view of the earlier noted high sensitivity of the sensor cells to certain compounds such as organic solvents and alcohols ^32^, iGIST bioassay followed a strict parallel design with every dose of PTX_#1_ evaluated against the matched dose of the PTX_#1_ solvent (SolC_#1_; 50% glycerol in H_2_O with 50 mM Tris, 10 mM glycine and 0.5 M NaCl). iGIST luminescence readout was firstly plotted as raw signals *vs* time (**Figure 2A-C**), which allows for quick visual assessment of the effects. Then, to obtain quantitative observer-independent estimate of the effects, we rendered the raw luminescence signals into numerical area under the curve (AUC) values, normalized to AUC of FSK response in the control sensor cells (not exposed to PTX_#1_ or SolC_#1_ before FSK stimulation). FSK response in the control cells served as an internal calibrator in the assay and was taken for 100% for every given run. The derived values were denoted as AUC%- values and utilized for deduction of PTX effects on Gαi signaling through pair-wise PTX_#1_ *vs* SolC_#1_ comparisons (**Figures 2D-E and S2**). Finally, to characterize Gαi signaling across a range of PTX_#1_ and SolC_#1_ exposures, we calculated the Gαi signal relay index (Gαi-SRI), expressed as a ratio of AUC%-values for FSK *vs* combination of FSK + Oct (AUC% _FSK_ / AUC% _FSK + Oct 10 nM_; **Figure 2F-H**) at every given PTX_#1_ and SolC_#1_ dose. At full abrogation of Gαi signaling by PTX, the sensor cells are expected to lose responsiveness to Oct, with Gαi-SRI approaching 1.0. Schematics of iGIST output values and of their calculations is shown in **Figure S1**.

**Figure 2.**
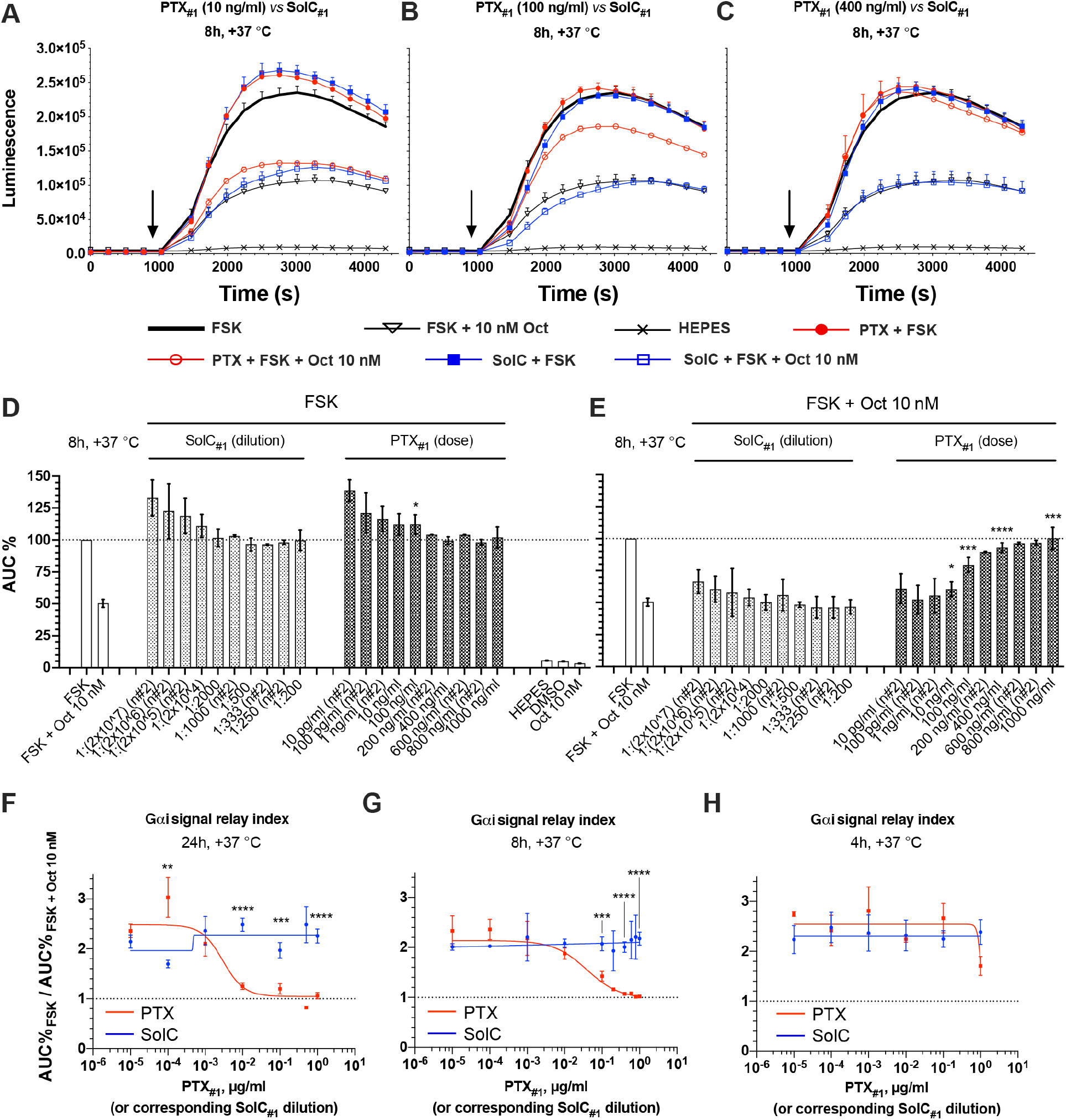
iGIST detects PTX_#1_-induced abrogation of Gαi signaling. **A-E)** FSK and Oct responses in the sensor cells after PTX_#1_ (w/v dose) or matched SolC_#1_ (corresponding stock dilution) exposure for 8h at +37 °C. Luminescence signals from a single representative experiment with the selected doses (A-C) and integrated results (as AUC%-values) of several independent runs (D-E). For curves in A-C, depicting raw luminescence reads, error bars denote +/- SD (only upper half shown) and y- and x-axes denote luminescence signal (AU) and time (s), respectively. The moment of FSK and Oct addition is indicated with the black arrow. For bar diagrams in D-E, y-axis depicts AUC%-values derived from the luminescence signals in several independent runs (response to FSK in control sensor cells that were not subjected to PTX_#1_ or SolC_#1_ is taken for 100%). Error bars represent average values +/- SEM. **F-H)** Oct/SSTR2-mediated effects on cAMP levels in sensor cells, measured as Gαi signal relay index (AUC% _FSK_ / AUC% _FSK + Oct 10 nM_ at a given dose of PTX_#1_ or SolC_#1_) after exposure to PTX_#1_ or matched SolC for 24, 8 and 4 h at +37 °C, respectively. Panels F-G depict integrated results of several independent runs (average values +/- SEM; see also **Figure S2C-D**). Panel H is based on data from a single representative experiment in 3x technical replicates (mean +/- SD; refer also to **Figure S2A-B**). Dose-response curves were fitted with a non-linear regression. A state of complete abrogation of Gαi signaling (AUC% _FSK_ / AUC% _FSK + Oct 10 nM_ = 1) is indicated with the black dotted line. All the assays were run in standard conditions, in 3x technical replicates. The number of individual assay repeats (n#) for bar diagrams ≥ 3, if not indicated otherwise. Significant differences for comparisons of responses at corresponding doses of PTX_#1_ *vs* SolC_#1_ are indicated with asterisks (further information in Experimental Section). Inferential statistics was only performed when n≥ 3 for individual assay repeats.

iGIST robustly registered PTX-induced abrogation of Gαi signaling that was proportional to the PTX_#1_ dose and time in contact with the cells. iGIST demonstrated the highest sensitivity at the longest PTX incubation studied (24 h), revealing a nearly complete abrogation of Gαi signaling at already 10 ng/ml of PTX (**Figures 2F and S2C-D)**. With shorter incubations, the PTX dose required for abrogation of Gαi signaling increased with the assay reliably capturing PTX_#1_ activity at 100 ng/ml with 8h incubation (**Figure 2D-E/G)**, and at around 1000 ng/ml with the 4h incubation (**Figures 2H and S2A-B)**. The global pattern of FSK response in PTX_#1_-treated sensor cells closely followed the one of SolC_#1_. Although comparisons of FSK responses at 100 ng/ml PTX_#1_ *vs* SolC_#1_ after 8h and 24h reached statistical significance (**Figures 2D and S2C**), the actual differences were very small, and thus likely had no practical relevance. The cAMP levels in the sensor cells without FSK stimulation were not significantly affected by PTX_#1_ across the dose range studied (**Figures 2A-C**, luminescent signals before black arrowhead), which is in line with the earlier reports ^27 29^.

As all the above evidence was obtained with a single PTX preparation (PTX_#1_), we validated the iGIST bioassay with another PTX formulation, from a different vendor and having a different solvent composition (PTX_#2_; in 10 mM Na_2_HPO_4_ and 50 mM NaCl in H_2_O). The response pattern of iGIST to PTX_#2_ was virtually identical to the one of PTX_#1_. Though the effect started to emerge at already 1 ng/ml of PTX_#2,_ abrogation of Gαi signaling became profound at 10 ng/ml of the toxin (**Figure 3B-C**) – the same threshold dose as with PTX_#1_ after 24h incubation. Apart from a borderline increase at 10 pg/ml, PTX_#2_ did not alter the pattern of FSK response, recapitulating the effects of SolC_#2_ (**Figure 3A**). Basal cAMP levels before FSK addition also stayed unaffected with PTX_#2_. Collectively, iGIST reliably detected PTX activity with two unrelated PTX preparations, revealing PTX-induced abrogation of Gαi signaling at ng/ml levels of the toxin.

**Figure 3.**
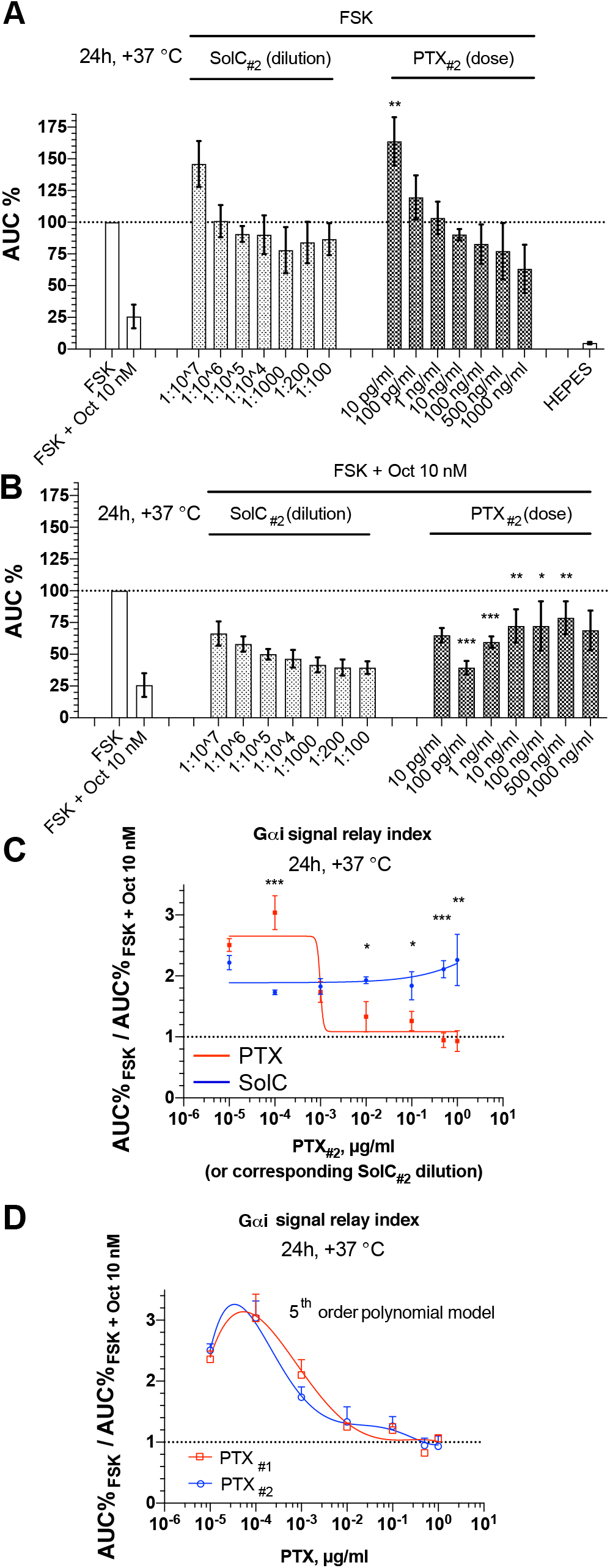
iGIST bioassay with PTX_*#2*_ and the biphasic model of PTX response. PTX_#2_ is a commercial PTX preparation from a different vendor and having a different solvent composition as compared to PTX_#1_. **A-B)** FSK and Oct responses in the sensor cells after PTX_#2_ or SolC_#2_ exposure for 24 h at +37 °C; integrated results of several independent runs (the number of individual assay repeats ≥ 3, with each assay in 3x technical replicates; shown are average values +/- SEM). The y-axis depicts AUC%-values derived from the luminescence signals (response to FSK in control sensor cells that were not subjected to PTX_#2_ or SolC_#2_ is taken for 100%). All the assays were run at standard conditions. Significant differences for comparisons of responses at corresponding doses of PTX_#2_ *vs* SolC_#2_ are indicated with asterisks (further info in Experimental Section). **C)** Oct/SSTR2-mediated effects on cAMP levels in the sensor cells, measured through Gαi signal relay index (i.e., ratio of AUC% _FSK_ and AUC% _FSK + Oct 10 nM_) after exposure to PTX_#2_ (w/v dose) or matched SolC (corresponding stock dilution) for 24 at +37 °C. Curves are based on the same data as shown in panels A-B (average values +/- SEM), and fitted through a non-linear regression (four-parameter logistic curve with a variable slope). The state of complete abrogation of Gαi signaling (AUC% _FSK_ / AUC% _FSK + Oct 10 nM_ ratio of 1.0) is depicted with the black dotted line. **D)** Gαi signal relay index (AUC% _FSK_ / AUC% _FSK + Oct 10 nM_) at different toxin doses, from PTX_#1_ and PTX_#2_ experiments with iGIST (same data points as on Figure 2F and Figure 3C); curve fitting through a 5^th^ order polynomial regression.

### iGIST reveals an unexpected potentiation of Gαi signaling at low PTX dose

When comparing PTX dose responses after 24 h, we unexpectedly detected a potentiation of Gαi signaling with 100 pg/ml of PTX, manifested as an increment of Gαi-SRI (**Figures 2F and 3C**). This effect seems paradoxical and difficult to understand from a standpoint of the canonical PTX activity, i.e. abrogation of Gαi signaling. Yet, the potentiation of Gαi signaling at 100 pg/ml PTX dose was highly reproducible, pronounced and consistently detected with both the toxin preparations (PTX_#1_ and PTX_#2_). Our initial model of PTX effect, based on Gαi-SRI and fitted through a non-linear regression (four-parameter logistic curve for an inhibitory response with a variable slope) could not accommodate this outlier. The resulting sigmoid curves (the red ones, **Figures 2F and 3C**) predicted a simple unidirectional inhibitory response from low-ng/ml levels of PTX onwards. The data urged us to consider an alternative model of PTX effect, which could be described by a bell-shaped curve with a truncated left arm when fitted through a 5^th^ order polynomial regression (**Figure 3D**). The resulting alternative model of PTX effects on Gαi signaling fits the experimental data much better. According to the alternative model, PTX exerts no effects on Gαi signaling at the lowest exposure tested (10 pg/ml), potentiates at 100 pg/ml dose and starts to abrogate at higher doses. Canonical abrogation of Gαi signaling with PTX is manifested first by a drop in Gαi-SRI back to the baseline level at around 1 ng/ml of the toxin (**Figure 3D**). This roughly corresponds to Gαi-SRI of 2 – the value reflective of Gαi signaling across the studied dose range of SolCs. Then, the effect continues to increase dose-dependently, reaching saturation with a nearly-complete abrogation of Gαi signaling at 10 ng/ml of PTX (Gαi-SRI of 1) (**Figure 3D**).

The potentiation of Gαi signaling by low-dose PTX in 24 h incubation, as revealed by the iGIST, is highly intriguing. Admittedly, the exact molecular basis remains a matter of subsequent studies. As for now, we hypothesize that the phenomenon relates to the dynamics of how the different G protein α-subunits are complexed and functionally regulated with G protein βγ-subunits ^38^. However, in view of the time scale of the potentiation effect, more complex compensatory mechanisms could be involved such as a transcriptional and/or translational response. Irrespectively of the nature of the underlying molecular mechanisms, detection of the potentiation effect has a profound application potential. First, it increases the sensitivity of iGIST two orders of magnitude, from *ca* 10 ng/ml down to 100 pg/ml of PTX. Secondly, it adds to the specificity of iGIST as reconstruction of the truncated bell-shape PTX response curve through serial dilution of an analyte would ensure the specific nature of the observed signal (**Figure 3D**). Collectively, iGIST reveals a hitherto undescribed potentiation effect of PTX on Gαi signaling, which improves specificity and increases the sensitivity of the iGIST to pg/ml range of PTX.

### iGIST bioassay is more sensitive than CHO cluster formation assay to detect PTX

Sensitivity of the iGIST bioassay was compared with that of the cluster formation assay (CFA), originally introduced by Hewlett et al. ^19^. To allow for direct comparison with iGIST bioassay, we obtained two strains of wild-type CHO cells from two different sources (designated as CHO_#1_ and CHO_#2_), and utilized the cells for time and dose-range studies with the earlier used toxin preparation (PTX_#1_). Through parallel use of the two CHO strains, representing the progeny of the same maternal CHO culture, we strived to mitigate the risks, associated with possible genetic and phenotypic drift in immortalized cell lines upon extended culturing ^39, 40^.

Both strains of CHO were subjected to live cell imaging with IncucyteHD imager under regular incubator conditions up to 72h from the moment of PTX addition. The derived phase-contrast images were visually graded by six independent observers, using a 3-tier scale (0 - no effect, 1 - ambiguous response, 2 - clear response; all comparisons - *vs* matched SolC) (**Figure 4**). With an arbitrary cut-off for a clear response set to 1.5, CHO_#1_ and CHO_#2_ demonstrated consistent and broadly similar performance in CFA. The lowest PTX_#1_ dose provoking a distinct phenotypic shift in both CHO strains was 10 ng/ml at 48 h. The perceived magnitude of phenotypic response increased further with PTX_#1_ dose, and both CHO_#1_ and CHO_#2_ generally served as reliable PTX sensors at PTX doses of ≥ 100 ng/ml. However, the exposure time required for PTX_#1_-induced morphological changes to emerge, did differ between the CHO strains. CHO_#2_ exhibited the pronounced phenotypic alteration only after 48-72 h of exposure, whilst CHO_#1_ underwent phenotypic switch earlier, already at 24 h, with morphological alterations becoming even more pronounced at 48 h. Shorter PTX_#1_ exposure times, i.e. 16 h or less, were insufficient for induction of clear phenotypic changes in either of the CHO strains even at the highest PTX_#1_ dose tested (500-1000 ng/ml).

**Figure 4.**
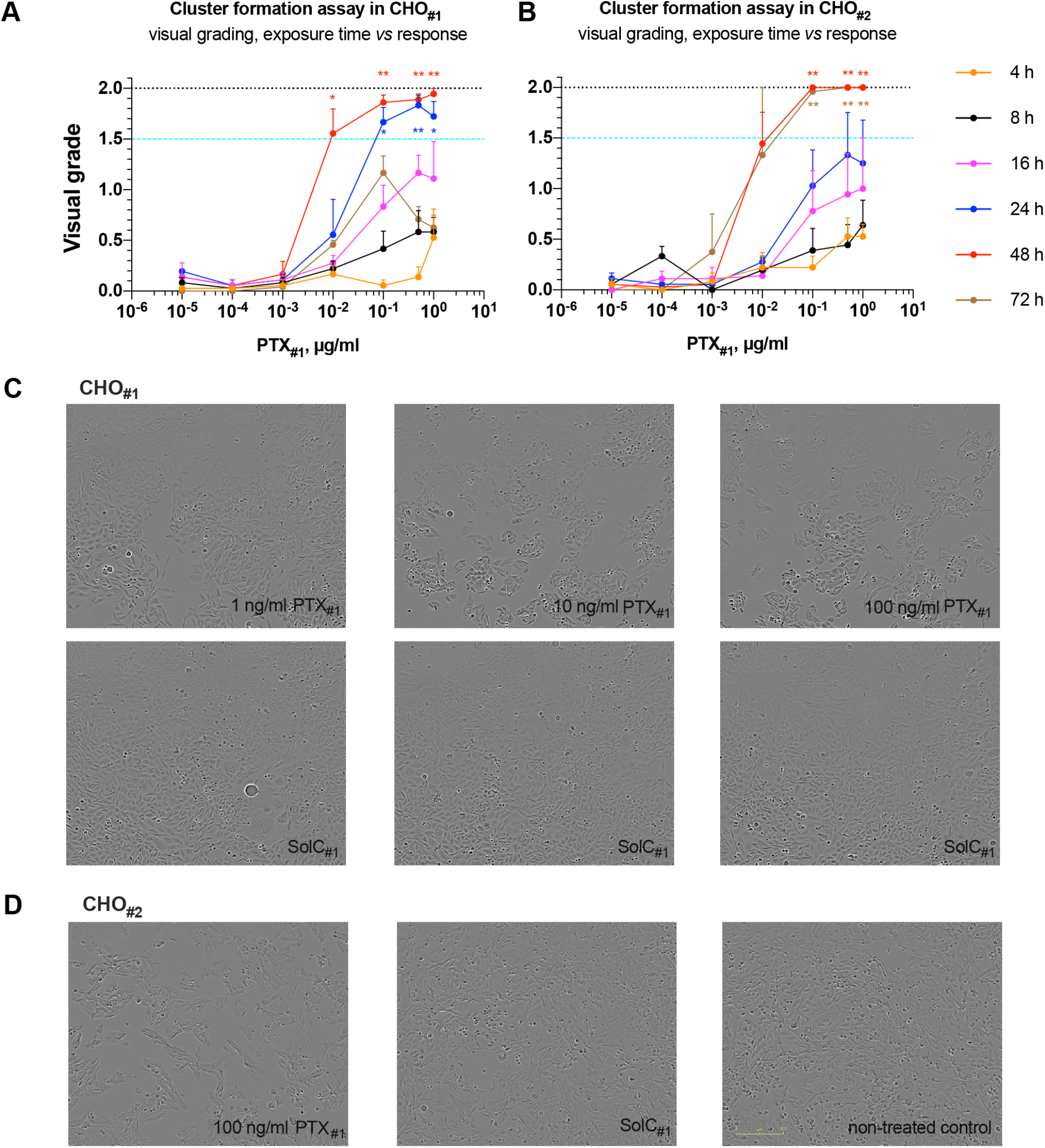
Cluster formation assay - visual grading of PTX_#1_-induced morphological changes in CHO cells. **A-B)** PTX_#1_ dose *vs* incubation time studies in two strains of CHO cells (CHO_#1_ and CHO_#2_), obtained from two different sources. CHO cells were seeded into 96-well plates, treated with the indicated doses of PTX_#1_ and continuously imaged with IncucyteHD under regular incubator conditions up to 72h. The resulting phase-contrast images were visually graded by independent observers. The y-axis depicts visual grade of morphological response in CHO (AU; average +/− SEM, with only upper half of SEM shown). The x -axis indicates the PTX dose. The maximal possible grade (2.0) and the pre-selected cut-off for a clear response (1.5) are indicated with black dotted and turquoise dashed lines, respectively. PTX exposures with a lower limit of 90% CI ≥ 1.0 or 1.5 for an average visual grade are marked with a single or double asterisk, respectively. The curve of 72 h is based on 2x independent experiments. All the other exposure times represent combined results of 3x independent experiments (each in at least 2x technical replicates). **C-D)** Phase-contrast images from the above CFA grading set, exposure of 48 h at +37 °C in CHO_#1_ or CHO_#2_. Stated are concentrations of PTX_#1_. Solvent and non-treated controls received matched dilutions of the SolC_#1_ or just HEPES buffer, respectively. Scale bar – 200 μm. Time-lapse movies of the phase-contrast images for CHO_#1_ with 100 ng/ml of PTX_#1_, as well for matched SolC_#1_ control, are available as **Supplementary Videos 1-2**.

If we compare only absolutely clear phenotypic responses to PTX_#1_, our CFA exhibits very close performance to the CFA in the original work of Hewlett et al. ^19^. Also, in view of the reportedly high variation in CFA results even with the same PTX preparations across different laboratories ^20, 41, 42^, the described results signify the robustness of our CFA. Of interest, automatic confluence analysis (IncuCyte software) of the same image set demonstrated a slightly improved resolution of PTX_#1_-induced effects in CHO as compared to the visual grading by the human observers (**Figure S3**). Here, a minor decline in estimated confluence was already noticeable at 1 ng/ml of PTX_#1_ at 48 h of treatment, with the effect becoming clear and pronounced from 10 ng/ml of PTX_#1_ onwards. The data underlines the subjective nature of visual grading in the conventional CFA, suggesting that observer-independent software-driven image analysis might make a better option for CFA. Most importantly, however, the data demonstrates that the iGIST bioassay is more sensitive to detect PTX_#1_ than CFA (*ca* 100-fold, with a threshold of 100 pg/ml of PTX) (**Figures 2F and S2C-D**).

### iGIST bioassay detects PTX spiked into Boostrix pertussis acellular vaccine

The commonly acknowledged limitation of CFA ^19^ and other proposed animal-free bioassays to detect PTX ^27, 29^ is their poor compatibility with the final vaccine product due to cytotoxicity of the aluminum-based adjuvants ^17^. This problem is pending, despite that several approaches to mitigate adjuvant toxicity, e.g. by means of vaccine dilution or barrier methods such as semi-permeable transwell inserts for culture plates, have been proposed ^17^. To analyze the applicability of iGIST for PTX detection in complex samples, i.e. commercial pertussis vaccines, we prepared ACV dilution series supplemented with a fixed PTX concentration. As industry-grade PTX-toxoid was not available, we spiked a known dose of the active PTX_#1_ to achieve a final concentration of 100 ng/ml into serial dilutions of PTX-toxoid-containing vaccine (Boostrix; includes 16 μg/ml of PTX-toxoid, admixed with tetanus and diphtheria toxoids). Despite the complexity of Boostrix, including significant levels of aluminum (≤ 0.08% w/v), detergent (Tween80) and two other toxoids (diphtheria and tetanus) with possible residual activity, iGIST successfully detected the spiked PTX_#1_. This was evidenced by the complete abrogation of Gαi signaling with Gαi-SRI of 1 in all the analyzed Boostrix dilutions (*≥*1:10) (**Figure 5**). The unspiked Boostrix was not neutral in terms of its effects on Gαi signaling in iGIST (**Figure 5C**, blue dots), but in the absence of the matched SolC the nature of the observed responses remains unknown. Importantly, we did not observe overt cytotoxicity (i.e., cell detachment, cell death) even at the most concentrated Boostrix solution tested (1:10 dilution) within the time window of the assay (24h) (**Figure S4**). Taken together, our results with the PTX_#1_-spiked Boostrix underline functional robustness of iGIST and highlight iGIST as a promising tool for PTX detection in complex samples.

**Figure 5.**
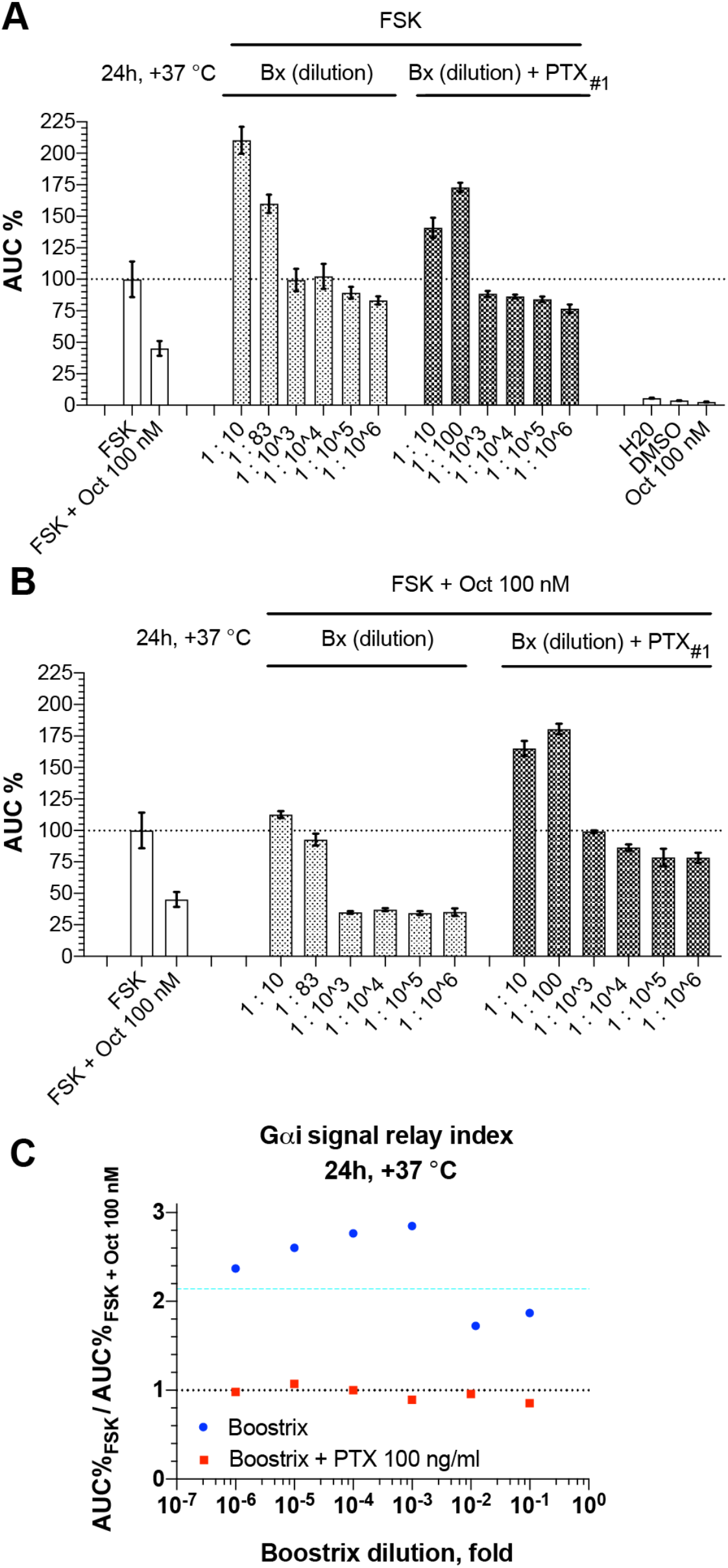
iGIST bioassay detects PTX_#1_ spiked into Boostrix pertussis ACV. **A-B**) FSK and Oct responses in the sensor cells after 24 h/+37 °C pre-incubation with serial dilutions of either Boostrix vaccine or Boostrix vaccine admixed with a fixed dose of PTX_#1_ (final PTX_#1_ concentration - 100 ng/ml). Data from a representative run in 3x technical replicates (average values +/− SD), performed under standard conditions. The y-axis indicates AUC%-values (response to FSK in control cells, received neither Boostrix nor Boostrix with PTX_#1_, is taken for 100%), Boostrix dilution of 1:83 on panels A-B (*vs* the intended 1:100 dilution) is a result of inadvertent pipetting mistake. **C**) Gαi signal relay index (AUC% _FSK_ / AUC% _FSK + Oct 10 nM_) *vs* vaccine dose, based on the data on panels A-B (average values depicted). Gαi signaling after exposure to SolC_#1_ and the state of complete abrogation of Gαi signaling are shown with turquoise dashed (Gαi-SRI = 2.14) and black dotted (Gαi-SRI = 1.0) lines, respectively. Gαi-SRI for 100 ng/ml of PTX_#1_ w/o vaccine equaled 1,022 (not shown). Higher Oct dose (100 nM) was utilized in the assay to ensure potent SSTR2 activation, thus minimizing possible effects of vaccine components on Gαi signaling.

### iGIST - objective digital readout and prospects for automation

An important advantage of iGIST is its observer-independency. This feature, combined with the microtiter plate format and the digital nature of iGIST readout, opens avenues for automatization, e.g. by means of robotic platforms for plate handling and luminescence acquisition. Data processing at a higher throughput might be approached through utilization of tailored scripts, rendering iGIST luminescence signals into numerical values, such as Gαi-SRI (AUC% _FSK_ / AUC% _FSK + Oct 10 nM_). For an alternative numerical index of PTX activity that streamlines data interpretation and thus might be more compatible with automated data processing, we propose a comparative Gαi-SRI, calculated as a ratio of (AUC% _FSK_ / AUC% _FSK + Oct 10 nM_) values for PTX-exposed and matched SolC-exposed samples (**Figure S5**). Reflective of the relative change in Gαi signaling in the sensor cells, be it potentiation or abrogation, and accounting for the effects of SolC, comparative Gαi-SRI should readily highlight PTX exposures.

Comparative Gαi-SRI can also be used to measure iGIST inter-assay variability, i.e. we obtained a 3-point composite estimate of 15,23% (equaling average coefficient of variation for comparative Gαi-SRIs for PTX 10 pg/ml, 100 pg/ml and 10 ng/ml at 24h of exposure, taken for no-effect level, maximal stimulation and inhibition, respectively; for 4x independent runs). Subsequent studies with appropriate controls, i.e. individual vaccine components and SolCs from different steps of PTX vaccine manufacturing process, are required to delineate the industry-scale applicability of iGIST.

## CONCLUSIONS

We established iGIST (<u>I</u>nterference in <u>G</u>αi-mediated <u>S</u>ignal <u>T</u>ransduction), a kinetic microtiter plate format bioassay to detect PTX at pg/ml levels by measuring its effect on inhibitory GPCR signaling. iGIST is observer-independent, has an objective digital readout and exceeds in sensitivity by 100-fold the currently used *in vitro* end-point technique to detect PTX activity, the cluster formation assay in Chinese hamster ovary cells ^17^. iGIST also detects PTX in complex samples, i.e. a commercial PTX-toxoid containing pertussis vaccine Boostrix that was spiked with an active PTX. We conclude that iGIST is a useful new tool for PTX basic research ^5^, PTX-targeted drug development ^43^, and PTX-focused industrial applications including the development and manufacturing of PTX-toxoid containing pertussis vaccines ^17^. Performed in microtiter plates, iGIST has potential for automation and batch processing, both required for industrial applications. Most importantly, iGIST is an animal-free assay and thereby emerges as promising alternative to complement or to replace mouse histamine sensitization test (HIST), the current vaccine industry standard to detect PXT activity ^17^. A better understanding of practical prospects for iGIST in industrial applications would follow from future rigorous head-to-head studies (iGIST *vs* comparators) with reference PTX and PTX-toxoid formulations, including the currently marketed pertussis vaccines.

## SUPPLEMENTARY MATERIAL

### Supplementary Figures 1-5

Figure S1. Schematic representation of iGIST output values and of their calculations.

Figure S2. iGIST AUC%-value results with 4h and 24h PTX_#1_ exposures.

Figure S3. Quantitation of PTX_#1_-induced clustering in monolayers of CHO cells by computer-aided confluence analysis.

Figure S4. Phase-contrast images of the sensor cells upon exposure to varying levels of Boostrix.

Figure S5. iGIST - objective digital readout and prospects for automation.

### Supplementary Videos 1-2

Video S1-S2. Time-lapse movies of the phase-contrast images for CHO_#1_ with 100 ng/ml PTX_#1_, aswell as with the matched SolC control (.mpg format videos 1-2, respectively).

## AUTHOR CONTRIBUTIONS

The manuscript was written through contributions of all authors. All authors have given approval to the final version of the manuscript. The authors declare that no competing interest exists.

## ACKNOWLEDGEMENTS

We thank Alexander V. Travov for excellent technical assistance (writing a script for processing of luminescence data) and critical outlook. The work was supported by European Community Mobility Programme EMA2 (grant #372117-1-2012-1-FI-ERAMUNDUS-EMA21; to VMP), Turku Doctoral Programme of Molecular Medicine (TuDMM; to VMP), K. Albin Johanssons Stiftelse (to VMP), Ida Montinin Säätiö (to VMP), Pentti and Tyyni Ekbom Foundation (to VMP), Instrumentarium Sciecne Foundation (to VMP); Sigrid Juselius Foundation (to ATP), Academy of Finland (grant no. 295296; to ATP).

## ABBREVIATIONS

AC/ACs: adenylyl cyclase/s
ACV/ACVs: acellular vaccine/s
ATCC: American Type Culture Collection
AU: arbitrary units
AUC: area under curve
cAMP: 3’5’-cyclic adenosine monophosphate
BSA: bovine serum albumin
CFA: cluster formation assay
CHO: Chinese hamster ovary cells
CI: confidence interval
CRE: cAMP response element
DMEM: Dulbecco’s Modified Eagle Medium
IndMed: inducing medium
FoV: field of view
FSK: forskolin
GPCR/GPCRs: G protein-coupled receptor/s
HIST: mouse histamine sensitization test
IC50: concentration of a compound, triggering half of the maximum inhibitory effect
iGIST: interference in Gαi-mediated signal transduction, a bioassay for PTX
iFBS: heat-inactivated fetal bovine serum
ns: non-significant
Oct: octreotide
ON: overnight
PTX: pertussis toxin
RT: room temperature
SD: standard deviation
SEM: standard error of the mean
SolC: solvent control (for PTX)
SRI: signal relay index
SSTR2: somatostatin receptor 2

## TABLE OF CONTENTS GRAPHIC

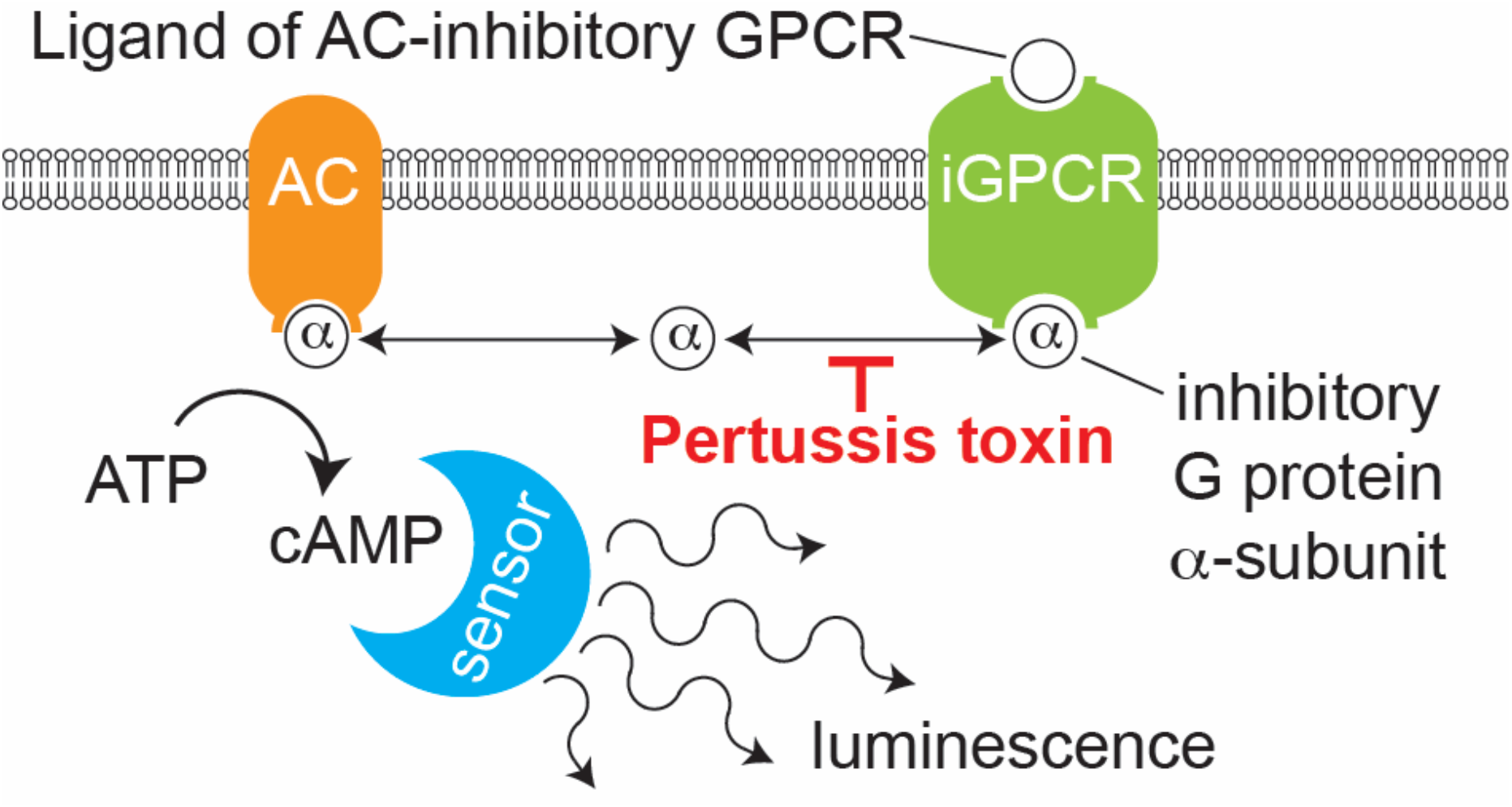

## Notes

### Competing Interest Statement

The authors have declared no competing interest.

