## supplemental material for "iGIST - a kinetic bioassay for pertussis toxin based on its effect on inhibitory GPCR signaling"

**SUPPLEMENTARY MATERIAL**

**Supplementary Figures 1-5**

Figure S1. Schematic representation of iGIST output values and of their calculations.

Figure S2. iGIST AUC%-value results with 4h and 24h $PTX_{\#1}$ exposures.

Figure S3. Quantitation of $PTX_{\#1}$-induced clustering in monolayers of CHO cells by computer-aided confluence analysis.

Figure S4. Phase-contrast images of the sensor cells upon exposure to varying levels of Boostrix.

Figure S5. iGIST - objective digital readout and prospects for automation.

**Supplementary Videos 1-2**

Video S1-S2. Time-lapse movies of the phase-contrast images for $CHO_{\#1}$ with 100 ng/ml $PTX_{\#1}$, as well as with the matched SolC control (.mpg format videos 1-2, respectively).

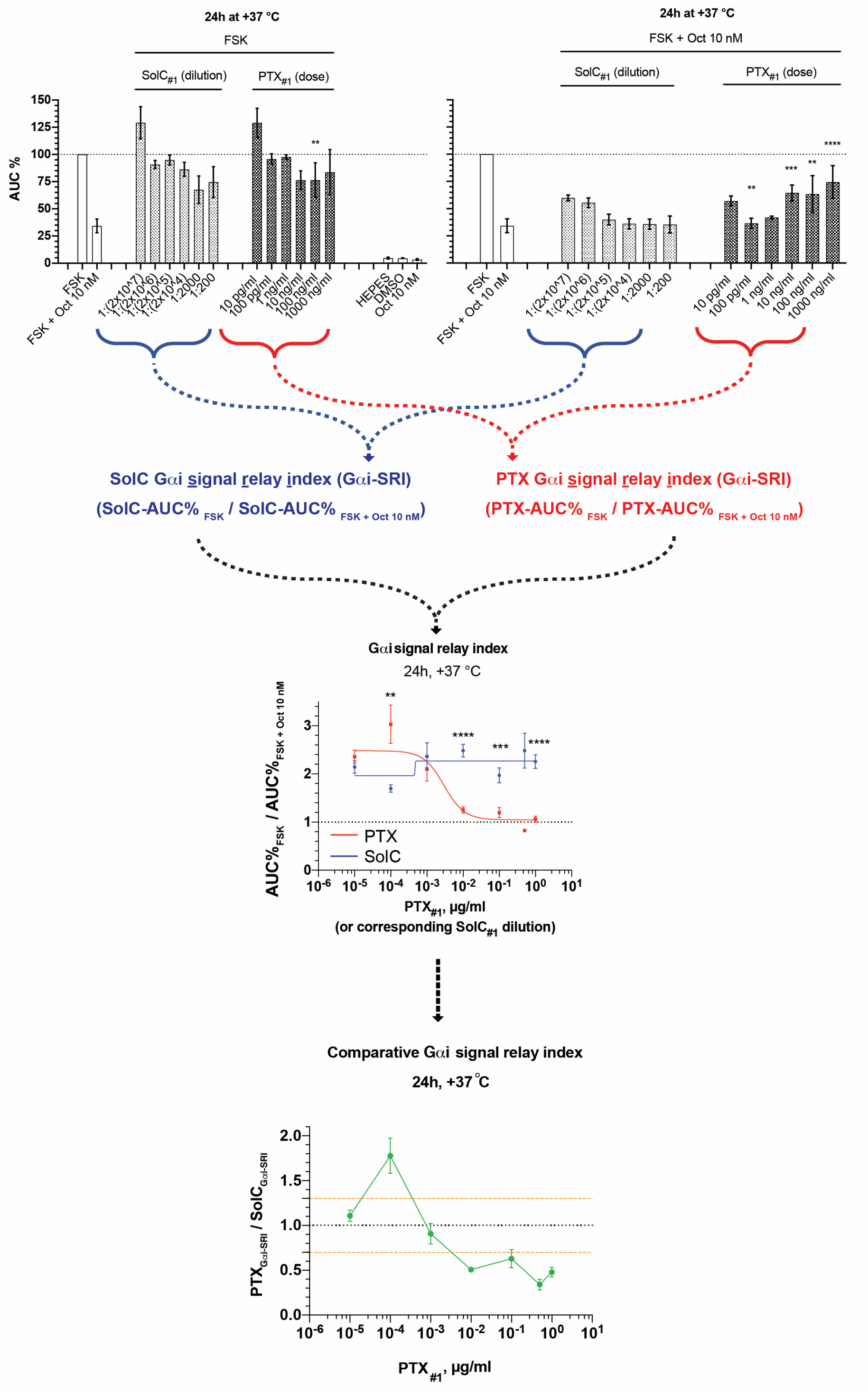


**Figure S1. Schematic representation of iGIST output values and of their calculations.** iGIST data is analyzed using three output values - **a)** AUC%-value (AUC-value normalized to AUC of FSK, which is set as 100% in each assay repeat); **b)** Gαi signal relay index (Gαi-SRI), equaling the ratio of $AUC\%_{FSK}$ to $AUC\%_{FSK + Oct\ 10\ nM}$ for PTX- or SolC-preincubated samples; **c)** Comparative Gαi signal relay index, equaling the ratio of Gαi-SRIs at matched SolC and PTX exposures [(SolC-$AUC\%_{FSK}$ / SolC-$AUC\%_{FSK + Oct\ 10\ nM}$) / (PTX-$AUC\%_{FSK}$ / PTX-$AUC\%_{FSK + Oct\ 10\ nM}$)]. For more information refer to the Experimental Section. The figure is based on data explained in detail in **Figures S2C-D**, **2F and S5.**

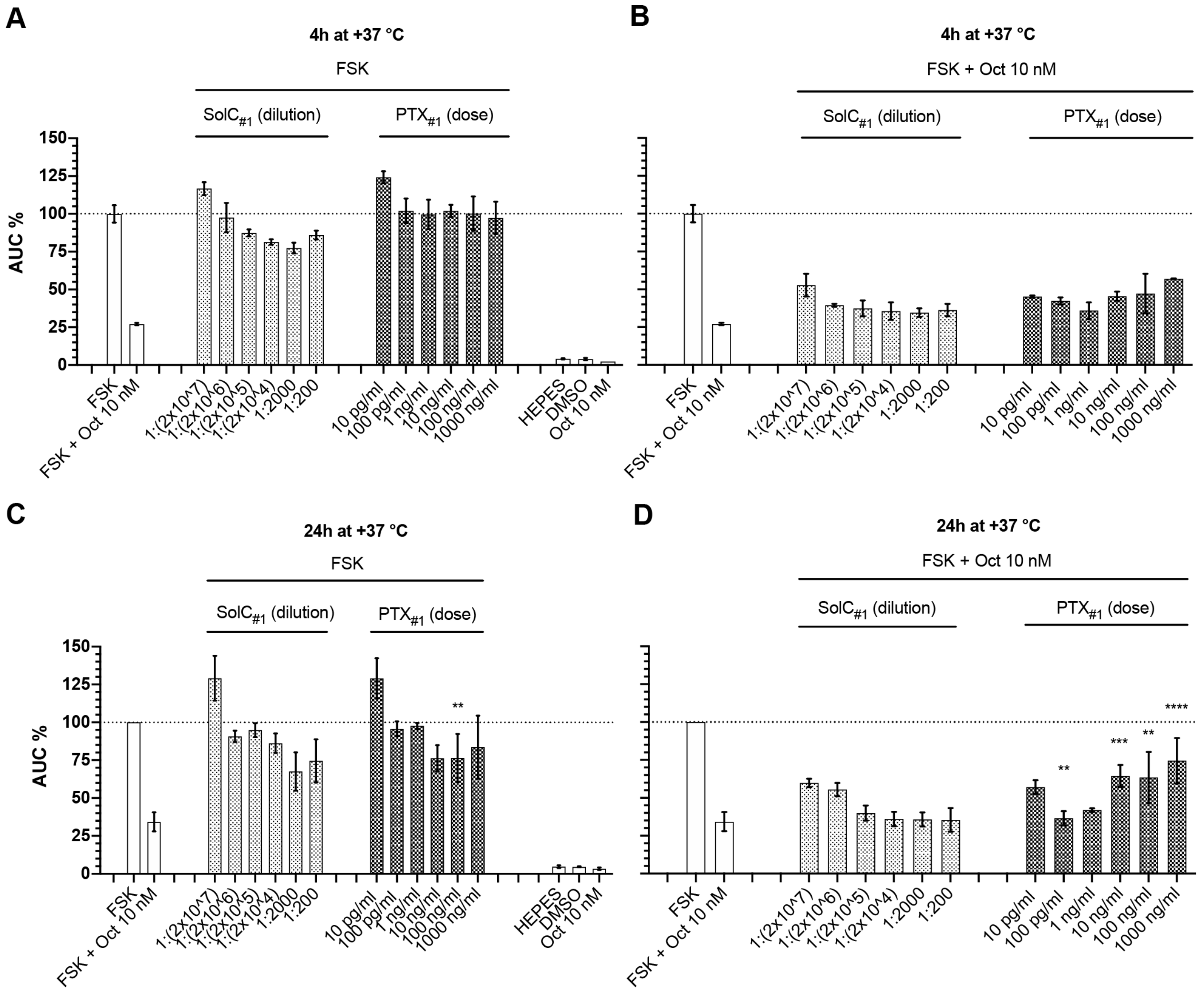


**Figure S2. iGIST AUC%-value results with 4h and 24h PTX#1 exposures. A-B/C-D**) FSK and Oct responses in the sensor cells after exposure to PTX#1 (w/v dose) or matched SolC#1 (corresponding stock dilution) for 4 h or 24 h at +37 °C. Data is shown from a single representative experiment (4 h) or several independent runs (24 h, the number of individual assay repeats ≥3). All the assays were run at standard conditions, in 3x technical replicates. The y-axis depicts AUC%-values (response to FSK in control sensor cells that were not exposed to PTX#1 or SolC#1 is taken for 100%), derived from the luminescence signals. Error bars for A-B and C-D represent average values +/-SD or +/- SEM, respectively. Significant differences for comparisons of responses at corresponding doses of PTX#1 *vs* SolC#1 at 24 h are indicated with asterisks (further info in Experimental Section). Abrogation of Gαi signaling, manifested as a decrease of Oct response in PTX#1-exposed samples (with corresponding rise in AUC%-values) as compared to SolC#1-exposed samples, becomes evident at 10 ng/ml with 24h of PTX exposure. The same effect could be even better perceived when luminescent responses are expressed in terms of Gαi-SRI (AUC% $_{FSK}$ / AUC% $_{FSK + 10 nM Oct}$) for PTX#1- *vs* SolC#1-exposed cells (refer to **Figure 2F**). iGIST also revealed an unexpected potentiation of Gαi signaling at low 100 pg/ml PTX dose with 24 h exposure (refer also to **Figure 2F and Figure 3C-D)**.

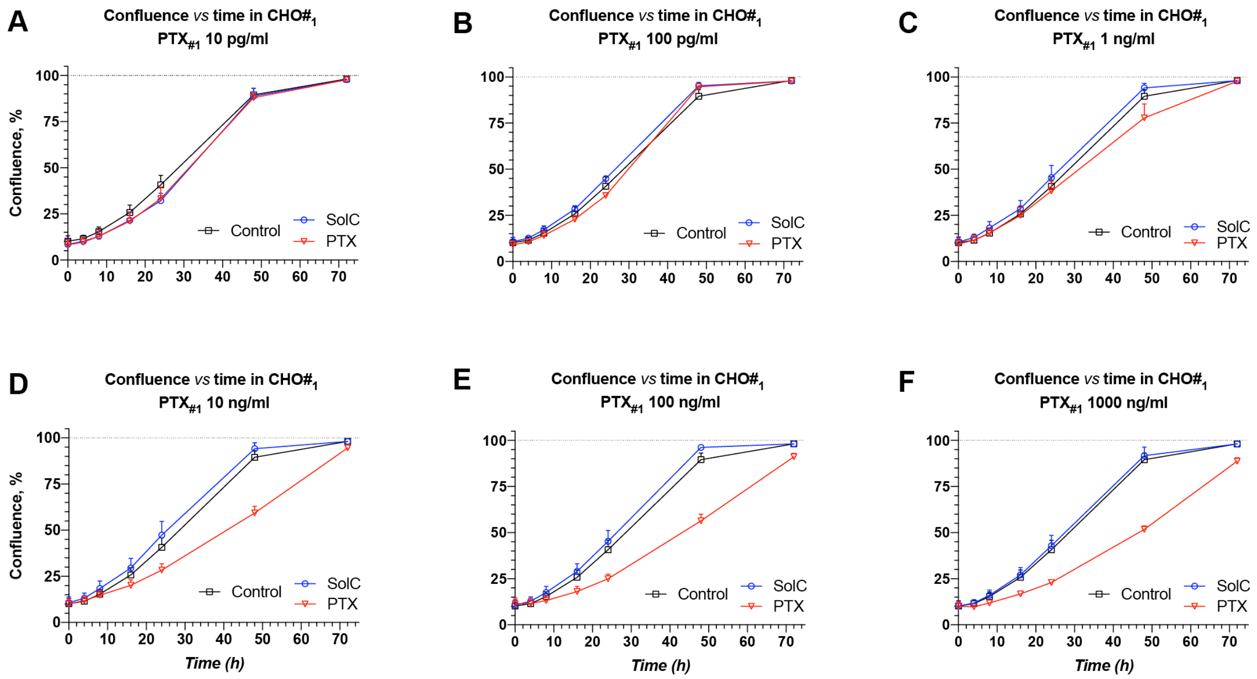


**Figure S3. Quantitation of PTX#1-induced clustering in monolayers of CHO cells by computer-aided confluence analysis. A-F**) Confluence charts for the indicated PTX#1 doses and incubation times in CHO#1, estimated with the in-built software package (Confluence v.1.5) of IncucyteHD on the same sets of phase-contrast images, employed for visual grading (**Figure 4**). The y-axis depicts confluence (%; average value +/- SEM, with only upper half of SEM shown). The x-axis depicts exposure time (h). The maximal possible achieved confluence (100%) is indicated with a black dotted line. All time points, apart from 72 h (2x independent repeats), are based on 3x individual experiments in at least 3x technical replicates each. Confluence analysis in CHO#2 under the same experimental conditions gave similar results (data not shown).

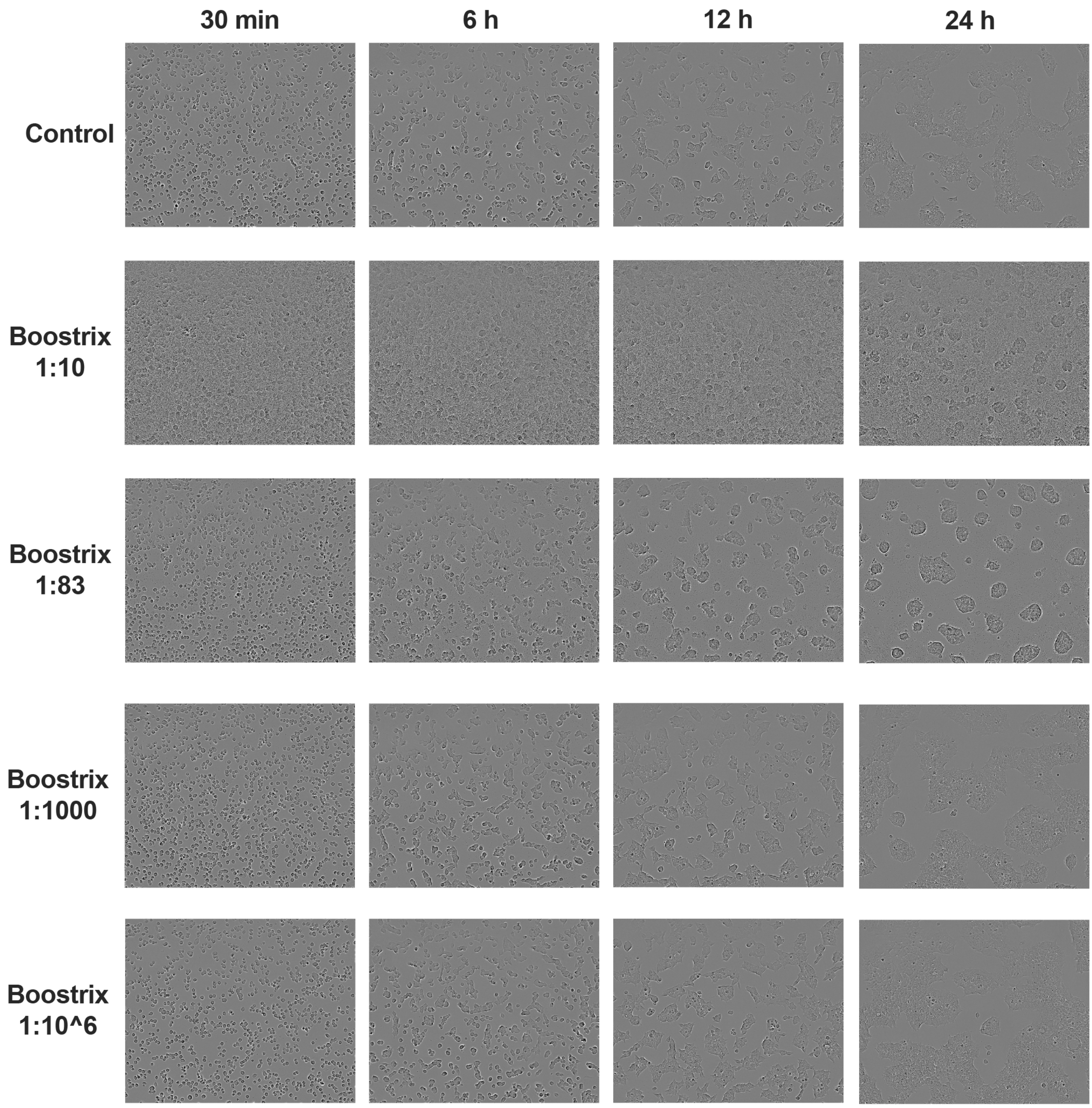


**Figure S4. Phase-contrast images of the sensor cells upon exposure to varying levels of Boostrix.** Imaging was performed with IncucyteHD, as described in the Experimental Section. Images were acquired within the same experiment described in Figure 5. Timing after Boostrix spiking is indicated. Sedimented vaccine particulate matter is clearly visible on the well surface and on the sensor cells at 1:10 and 1:83 dilutions. Control cells received matched volumes of $H_2O$. Boostrix was not neutral in terms of effects on cellular morphology at the highest exposure (1:10 and 1:83 dilutions), with phenotypic alterations becoming evident >6h after vaccine spiking. Yet, no gross cytotoxicity (cell detachment, cell death) was observed within the time span of the experiment.

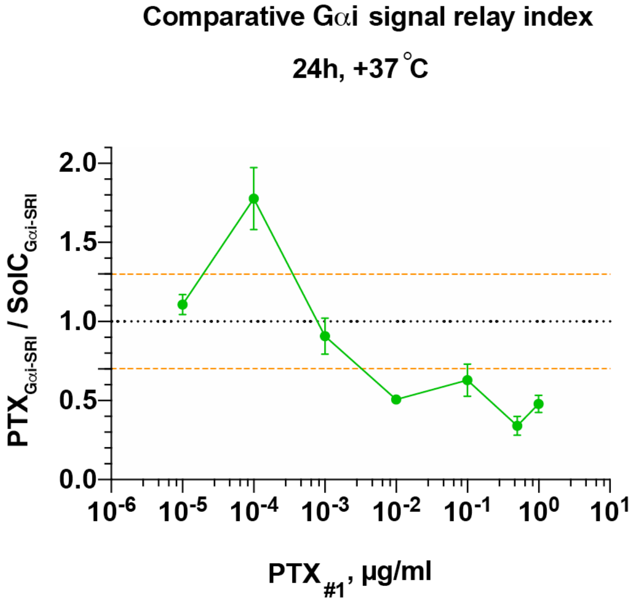


**Figure S5. iGIST - objective digital readout and prospects for automation.** The comparative Gαi signal relay index after 24 h PTX exposure. The index is calculated as a ratio of AUC% $_{FSK}$ / AUC% $_{FSK\ +\ Oct\ 10\ nM}$ values, i.e. Gαi-SRI values, for PTX$_{\#1}$ to matched SolC$_{\#1}$ from Figure 2F. The y-axis depicts average values +/- SEM). When combined with an arbitrary threshold of more than +/- 25% (depicted with orange dashed lines) from the no-effect level (1.0; black dotted line), the comparative Gαi-SRI readily highlights PTX dose of 100 pg/ml as potentiating Gαi signaling and PTX doses of 10 ng/ml and higher as abrogating Gαi signaling. The comparative Gαi-SRI, encompassing the effects of PTX and SolC in a single numerical value and revealing alterations in Gαi signaling in any direction, should facilitate automated quantitative analysis of iGIST data upon assay up-scaling.

**Supplementary Videos 1-2**

Time-lapse movies of the phase-contrast images for $CHO_{\#1}$ with 100 ng/ml of $PTX_{\#1}$, as well as with the matched SolC control (.mpg format videos 1-2, respectively). Videos span a period of 72h of live cell imaging under incubator conditions with IncucyteHD. CHO cells were spiked with PTX or SolC control within 5 minutes before time point zero.
